# Corneal keratocytes, fibroblasts, and myofibroblasts exhibit distinct transcriptional profiles *in vitro*

**DOI:** 10.1101/2024.02.28.582620

**Authors:** Kara Poole, Krithika S. Iyer, David W. Schmidtke, W. Matthew Petroll, Victor D. Varner

## Abstract

**Purpose:** After stromal injury to the cornea, the release of growth factors and pro-inflammatory cytokines promotes the activation of quiescent keratocytes into a migratory fibroblast and/or fibrotic myofibroblast phenotype. Persistence of the myofibroblast phenotype can lead to corneal fibrosis and scarring, which are leading causes of blindness worldwide. This study aims to establish comprehensive transcriptional profiles for cultured corneal keratocytes, fibroblasts, and myofibroblasts to gain insights into the mechanisms through which these phenotypic changes occur.

**Methods:** Primary rabbit corneal keratocytes were cultured in either defined serum-free media (SF), fetal bovine serum (FBS) containing media, or in the presence of TGF-β1 to induce keratocyte, fibroblast, or myofibroblast phenotypes, respectively. Bulk RNA sequencing followed by bioinformatic analyses was performed to identify significant differentially expressed genes (DEGs) and enriched biological pathways for each phenotype.

**Results:** Genes commonly associated with keratocytes, fibroblasts, or myofibroblasts showed high relative expression in SF, FBS, or TGF-β1 culture conditions, respectively. Differential expression and functional analyses revealed novel DEGs for each cell type, as well as enriched pathways indicative of differences in proliferation, apoptosis, extracellular matrix (ECM) synthesis, cell-ECM interactions, cytokine signaling, and cell mechanics.

**Conclusions:** Overall, these data demonstrate distinct transcriptional differences among cultured corneal keratocytes, fibroblasts, and myofibroblasts. We have identified genes and signaling pathways that may play important roles in keratocyte differentiation, including many related to mechanotransduction and ECM biology. Our findings have revealed novel molecular markers for each cell type, as well as possible targets for modulating cell behavior and promoting physiological corneal wound healing.

## INTRODUCTION

The cornea is the soft tissue at the anterior aspect of the eye that is responsible for bending light towards the retina. It consists of three primary cellular layers: the epithelium, stroma, and endothelium, with the stroma accounting for 90% of the corneal thickness^1–3^. Throughout this stromal layer, corneal keratocytes^3, 4^ are embedded within a highly organized extracellular matrix (ECM), consisting primarily of lamellae of aligned collagen fibrils^1, 2, 5^. In their quiescent state, keratocytes maintain the structure of the ECM^6–8^, which is necessary for corneal transparency^9^. However, following injury, various growth factors, such as transforming growth factor beta 1 (TGF-β1), platelet derived growth factor-BB (PDGF-BB), and fibroblast growth factor (FGF), among others, enter the stromal space and initiate a wound healing response. During this process, corneal keratocytes transform into a repair phenotype that involves differentiation into either fibroblasts or myofibroblasts^4, 10, 11^. Corneal fibroblasts are characterized by increased proliferation and migration towards the site of injury^4, 12^, whereas myofibroblasts adopt a contractile and fibrotic phenotype, in which they generate elevated traction forces, express alpha-smooth muscle actin (α-SMA), and secrete a variety of fibrotic ECM proteins^7, 13–21^. Myofibroblast differentiation is thought to be driven by signaling downstream of TGF-β1 and, in protracted cases, is associated with corneal haze and a loss of visual acuity^22, 23^.

To study these stromal cell phenotypes *in vitro*, three distinct culture models have been established experimentally: 1) primary corneal keratocytes cultured in defined serum-free (SF) media to maintain a quiescent phenotype, 2) keratocytes cultured with fetal bovine serum (FBS) to transform them into corneal fibroblasts, and 3) keratocytes cultured in the presence of TGF-β1 to induce myofibroblast differentiation^24, 25^. Using these culture conditions, previous studies have investigated the many phenotypic differences between keratocytes, fibroblasts, and myofibroblasts, with several focusing on differences in the morphology, motility, and mechanical behavior of these cells^19, 20, 26–28^. Many of these studies have highlighted the importance of cell-ECM interactions, indicating how the composition, structure, and mechanical properties of the ECM can regulate cell contractility, migration, and ECM remodeling^19, 24, 27, 29, 30^. For example, the TGF-β1-induced differentiation of myofibroblasts from quiescent corneal keratocytes is highly sensitive to changes in ECM stiffness^20, 24^. However, it is still unclear how these cells sense and transduce these biomechanical signals into changes in differentiation and behavior.

An investigation of the global transcriptional profile associated with each of these cell types could offer important information on the changes in gene expression associated with keratocyte differentiation into fibroblasts and myofibroblasts. Such an approach also offers the potential to pinpoint new target molecules within the signaling pathways that precede each of these phenotypic changes. Here, we used bulk RNA sequencing (RNA-seq) to establish comprehensive transcriptional profiles for cultured corneal keratocytes (SF), fibroblasts (FBS), and myofibroblasts (TGF-β1). Differential gene expression and functional analyses provided new insight into the mechanisms that regulate each of these cell phenotypes. Of particular interest, we identified several differentially expressed genes and pathways related to cell biomechanics and ECM remodeling.

## METHODS

### Isolation and Cell Culture of Primary Rabbit Keratocytes

Normal rabbit keratocytes (NRKs) were isolated from New Zealand white rabbit eyes (Pel-Freez Biologicals, Rogers, AR) and cultured as described previously^25^. Briefly, after removing the epithelium and endothelium, dissected corneal buttons were digested overnight at 37°C in culture media containing 2.0 mg/mL collagenase (Gibco, Grand Island, NY) and 0.5 mg/mL hyaluronidase (Worthington Biochemical Corporation, Lakewood, NJ). The isolated cells were then centrifuge-pelleted, resuspended in defined SF media containing Dulbecco’s modified Eagle’s medium (DMEM; Sigma-Aldrich, St. Louis, MO) supplemented with 100 µM non-essential amino acids (Gibco, Grand Island, NY), 100 µg/mL ascorbic acid (Sigma-Aldrich, St. Louis, MO), 1% RPMI vitamins solution (Sigma-Aldrich, St. Louis, MO), and 1% antibiotic antimycotic solution (Sigma-Aldrich, St. Louis, MO), and plated in 25 cm^2^ tissue culture flasks. After 4 days of SF culture, first passage NRKs were plated at a density of 30,000 cells/mL on glass coverslips coated with 50 µg/mL type I collagen (Advanced BioMatrix, Carlsbad, CA)^31^. The cells were allowed to attach to the substrates over 24 hr, then the culture media was swapped for either defined SF media, SF media supplemented with 5 ng/mL TGF-β1 (Sigma-Aldrich, St. Louis, MO), or DMEM containing 10% FBS (Sigma-Aldrich, St. Louis, MO). The cells were then cultured for 5 days (with a media swap at 48 hr) for both SF and TGF-β1 conditions or for 3 days following exposure to FBS to account for the increased proliferation rate in this condition.

### RNA Isolation and Sequencing

Eight samples of total RNA were collected for each culture condition using the Aurum Total RNA Mini Kit (Bio-Rad, Hercules, CA). Briefly, cells on collagen-coated glass coverslips were washed twice with sterile phosphate-buffered saline (PBS) and lysed using Aurum Total RNA Lysis Solution. The lysate was collected into sterile 1.5 mL tubes and mixed with RNase-Free 70% ethanol before being added to Aurum RNA-Binding Mini Columns. A series of centrifugation and low- and high-stringency solution washing steps were performed before eluting the RNA from the columns using molecular biology grade water. The concentration and purity of each sample were confirmed using a Thermo Scientific NanoDrop One^C^. For each culture condition, two experimental replicates (with two biological and two technical replicates for each experiment) were sent to Novogene Co. (Sacramento, CA) for bulk RNA-seq on Illumina platforms.

### Bioinformatic Analysis

Initial bioinformatic analysis performed by Novogene included sample and data quality control, mapping to the reference genome (Hisat2 v2.0.5), gene expression quantification (featureCounts v1.5.0-p3), differential expression analysis (DESeq2 R package v1.20.0), and enrichment analysis (clusterProfiler R package v3.8.1). The reference genome used for sequence mapping was generated by the McDermott Center Next Generation Sequencing Core at UT Southwestern Medical Center (Dallas, TX).

#### Gene Expression Quantification

FPKM (Fragments Per Kilobase of transcript per Million fragments mapped) values were calculated based on the length and read count mapped to each gene. FPKM considers the effects of both sequencing depth and gene length and is currently the most common method for estimating gene expression levels. Principal component analysis (PCA) was performed in MATLAB using FPKM values to compare the clustering of samples and culture conditions. PCA is the method of algebraically reducing the dimensionality and extracting significant components from several gene variables to evaluate differences within a group and between different groups in a set of samples^32^.

#### Differential Expression Analysis

Differential expression analysis was performed for the following comparisons: FBS vs. SF, TGF-β1 vs. FBS, and TGF-β1 vs. SF. Parametric analysis of variance (ANOVA) with Benjamini-Hochberg False Discovery Rate correction at p = 0.05 was performed on normalized data to identify genes that were significantly differentially expressed between groups (adjusted p-value ≤ 0.05 and absolute value of log2(fold change) ≥ 1). Heatmaps were created in GraphPad Prism using normalized, log2-transformed FPKM values (log2(FPKM+1)) to display relative expression levels across samples (columns) for sets of differentially expressed genes (rows). To normalize expression levels, each sample’s gene expression value was adjusted by subtracting the average expression of that gene across all samples, then dividing the result by the standard deviation of the gene’s expression. In addition, bar plots showing average FPKM values among the different culture conditions were generated for selected genes of interest.

#### Enrichment Analysis

Sets of significant differentially expressed genes were further analyzed through Kyoto Encyclopedia of Genes and Genomes (KEGG) enrichment analysis, and pathways with padj < 0.05 were considered significantly enriched. Bar plots were generated in GraphPad Prism to visualize pathway significance, the number of up- or downregulated genes associated with each pathway, and the BRITE functional hierarchies under which each pathway is classified. For a subset of significantly enriched KEGG pathways, volcano plots were generated in GraphPad Prism to visualize the significance and log2(fold change) of individual genes.

### Fluorescence Microscopy

In other experiments, samples were fixed and stained for fluorescence microscopy. Additional substrates for each culture condition were used for F-actin and nuclei labeling and immunostaining for fibronectin, vimentin, or α-SMA to observe cytoskeletal organization, morphological changes, and the presence of markers commonly associated with each phenotype. Cells were fixed in 3% paraformaldehyde in PBS for 10 min at room temperature, washed three times with PBS, then permeabilized in 0.5% Triton X-100 in PBS for 15 min. The samples were then blocked with 1% bovine serum albumin fraction V (Equitech-Bio, Kerrville, TX) in PBS for 1 hr at room temperature and incubated overnight at 4°C with the primary antibody solution. The following primary antibodies were used: anti-fibronectin (1:200 dilution; sc-18825; Santa Cruz Biotechnology, Dallas, TX), anti-vimentin (1:200 dilution; sc-6260; Santa Cruz Biotechnology, Dallas, TX), and anti-α-SMA (1:250 dilution; A5228; Sigma-Aldrich, St. Louis, MO). After washing three times, samples were then incubated with Alexa Fluor conjugated secondary antibody, as well as Alexa Fluor 546 phalloidin (Invitrogen, Waltham, MA) for 2 hr at room temperature, followed by 4’-6-diamidino-2-phenylindole (DAPI; Invitrogen, Waltham, MA) for 20 min. Imaging was performed on a Zeiss LSM 800 laser scanning confocal microscope using a 40×, NA 1.3, Oil DIC Plan-Apochromat objective.

### Western Blotting

In other experiments, after rinsing twice with ice-cold, sterile PBS, protein was collected from cells cultured in either SF, FBS, or TGF-β1 conditions using a lysis buffer solution containing Pierce RIPA Buffer and Halt Protease & Phosphatase Inhibitor Cocktail (Thermo Scientific, Waltham, MA). Sample lysates were mixed for 30 min at 4°C, then centrifuged at 10,000G for 10 min at 4°C. Protein concentrations were measured using the Microplate BCA Protein Assay Kit (Thermo Scientific, Waltham, MA) to determine load volumes for 5 µg of total protein. Protein samples were subjected to SDS-PAGE electrophoresis using Bio-Rad Mini-PROTEAN TGX Gels, then transferred to PVDF membranes (Bio-Rad, Hercules, CA). Membranes were stained for total protein using Ponceau S (Sigma-Aldrich, St. Louis, MO) and subsequently probed with primary antibody followed by an anti-mouse IgG HRP-linked antibody (Cell Signaling, Danvers, MA). The following primary antibodies were used: anti-aldehyde dehydrogenase 1-A1 (1:1000 dilution; sc-374149; Santa Cruz Biotechnology, Dallas, TX), anti-vimentin (1:1000 dilution; sc-6260; Santa Cruz Biotechnology, Dallas, TX), or anti-α-SMA (1:1000 dilution; A5228; Sigma-Aldrich, St. Louis, MO). Colorimetric (total protein) or chemiluminescence (target protein) imaging was performed using a GE Healthcare Amersham Imager 600 Series. In all cases, Image Studio software (version 5.2.5) was used to quantify protein expression and the amount of target protein was normalized to the amount of total protein. Statistical analysis was performed using GraphPad Prism 10. A one-way ANOVA with a Tukey post-hoc test was used to compare group means, and a p-value of less than 0.05 was considered statistically significant.

## RESULTS

### Distinct transcriptional profiles characterize corneal keratocytes in SF, FBS, and TGF-β1 culture conditions

We conducted bulk RNA-seq experiments using primary NRKs cultured in defined SF media, or in the presence of either FBS or exogenous TGF-β1. PCA showed distinct transcriptional differences between cells in each culture condition (Fig. 1A). Technical and experimental replicates clustered closely together in plots of PC1 vs. PC2, while the clusters associated with each culture condition were clearly separated. This result suggested the presence of a distinct gene expression profile for the cells within each group. Subsequent differential expression analysis revealed numerous DEGs among the different culture conditions (Fig. 1B). A comparison of FBS vs. SF contained the highest total number of DEGs at 7693, followed by TGF-β1 vs. FBS at 5704, and TGF-β1 vs. SF at 4430. DEGs present in more than one comparison were indicated by overlapping areas on the Venn diagram (Fig. 1B). Cumulatively, 3174 genes were differentially expressed if both FBS- and TGF-β1-treated cells were compared with SF, while 1311 genes were differentially expressed among all three groups.

**Figure 1.**
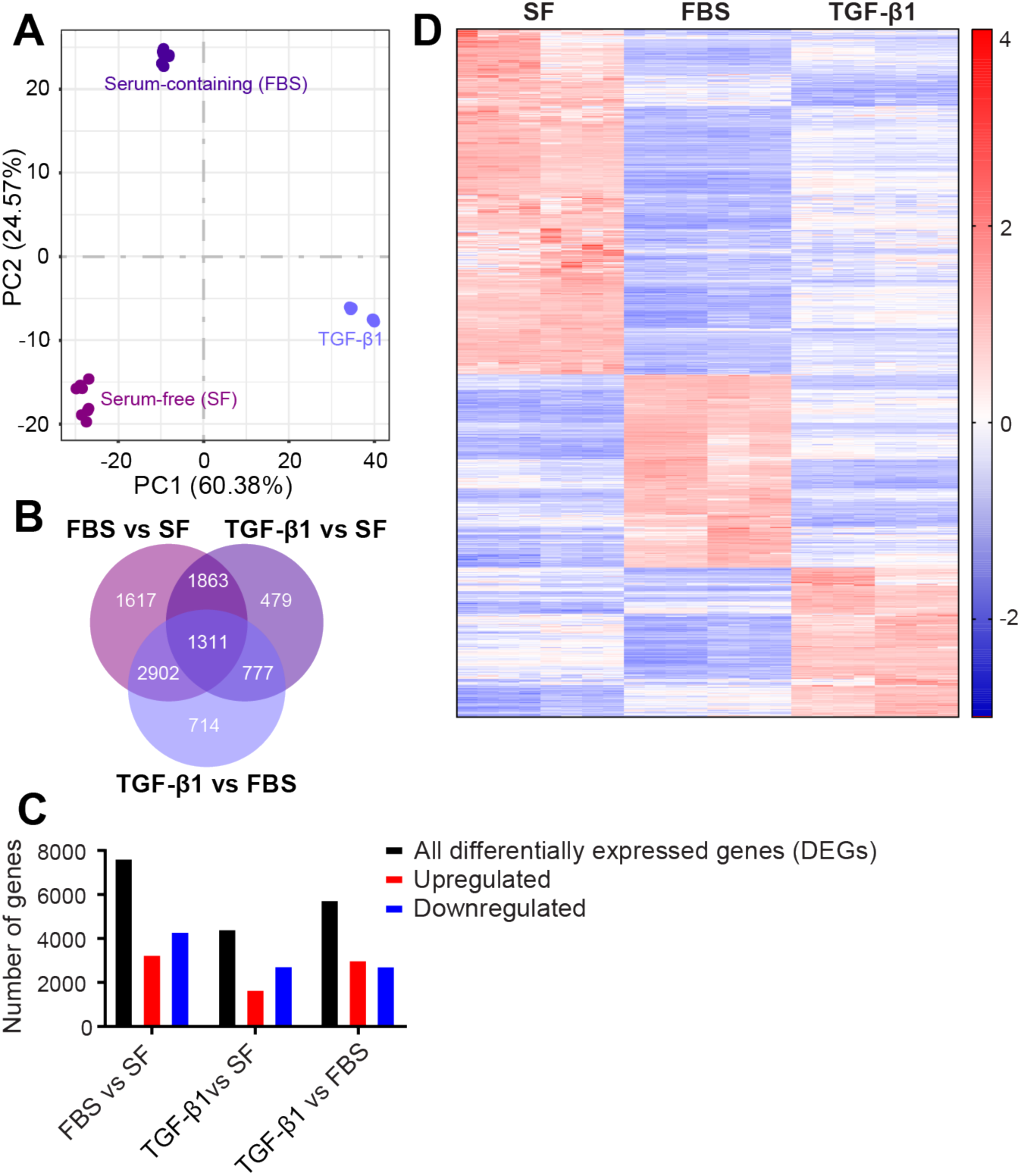
Distinct transcriptional profiles characterize corneal keratocytes in SF, FBS, and TGF-β1 culture conditions. (A) Principal component analysis (PCA) of transcriptional differences between corneal keratocytes cultured in defined serum-free (SF) media, FBS, or TGF-β1. Principal component (PC) 1 and PC2, account for 60.38% and 24.57% respectively, of the variability among these groups. Analysis was performed using normalized, log2-transformed FPKM values. (B) Venn diagram shows the number of significant differentially expressed genes (DEGs) in each comparison. DEGs with an adjusted p-value (padj) ≤ 0.05 and | log2(fold change) | ≥ 1 were considered significant. (C) Bar plot represents the number of DEGs either up- or downregulated in each comparison. (D) Heatmap of normalized, log2-transformed FPKM values (log2(FPKM+1)) for genes that were differentially expressed across all three culture conditions. Red color indicates genes with high relative expression levels and blue color indicates genes with lower expression levels.

We also observed distinct differences in the number of up- and downregulated DEGs when pairwise comparisons were made between culture conditions (Fig. 1C). A greater fraction of DEGs was downregulated when comparing either FBS or TGF-β1 to SF, whereas a slightly larger fraction of DEGs was upregulated when comparing TGF-β1 to FBS. A heatmap of DEGs among all groups indicated further that a distinct transcriptional profile was associated with each culture condition (Fig. 1D). Approximately 50% of these DEGs had high relative expression in SF, while 28% and 22% showed high expression in FBS and TGF-β1, respectively. Thus, the principal component, gene count, and differential expression analyses illustrate distinct and significant differential gene expression patterns between keratocytes cultured in SF media, FBS or TGF-β1.

### Markers associated with keratocytes, fibroblasts, and myofibroblasts are differentially expressed between culture conditions

We then compared our transcriptional data with molecular markers known to be associated with corneal keratocytes, fibroblasts, and myofibroblasts. Proteoglycans and corneal crystallins known to be important for corneal transparency^2, 33, 34^, such as keratocan (KERA), mimecan (OGN), decorin (DCN), lumican (LUM), aldehyde dehydrogenase 1 family member A1 (ALDH1A1), and transketolase (TKT), showed high relative expression in SF culture (Fig. 2A). In a consistent manner, analysis of western blot data confirmed elevated ALDH1A1 levels when comparing SF to either FBS or TGF-β1, indicating a quiescent keratocyte phenotype in this culture condition (Fig. 2B).

**Figure 2.**
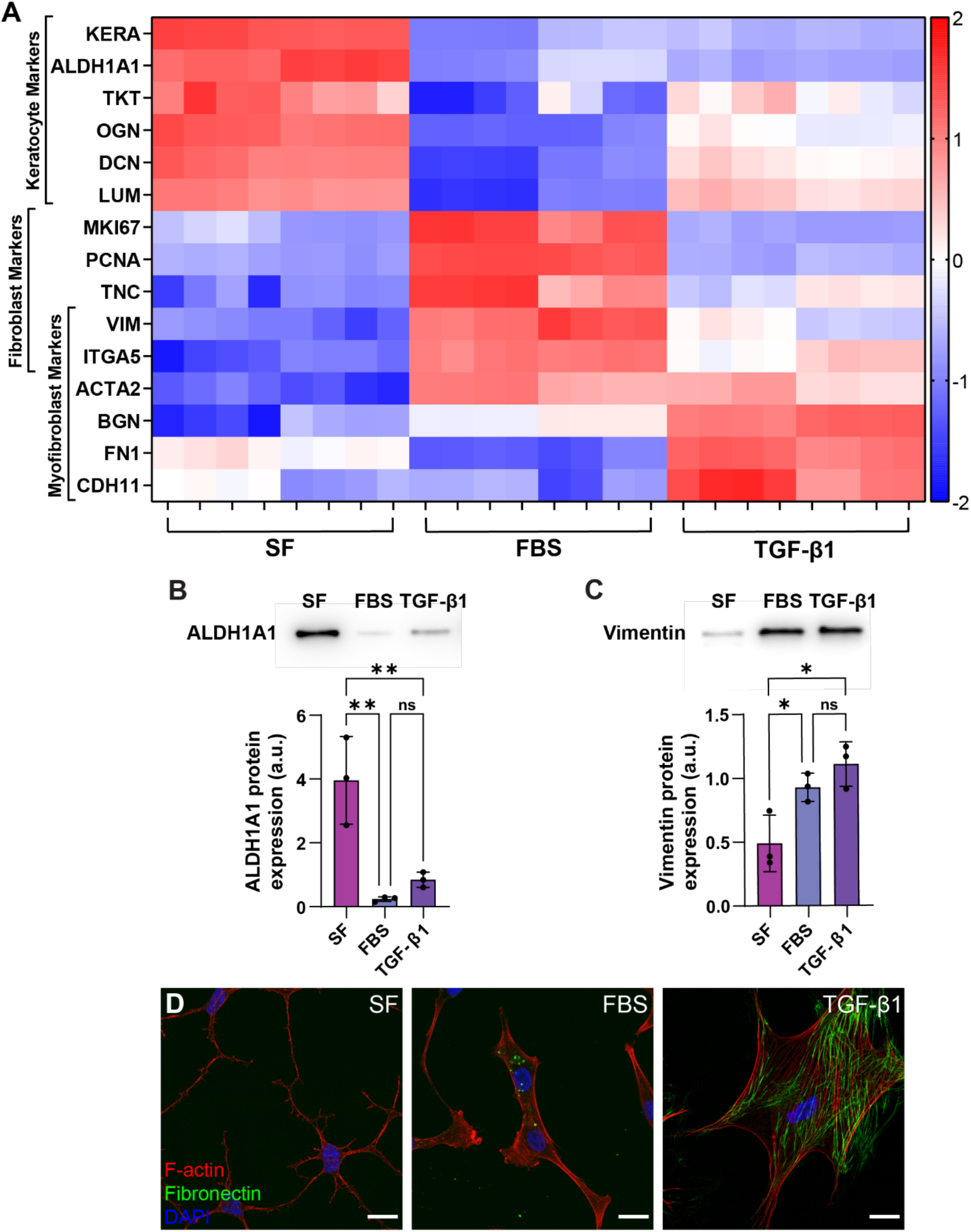
Markers associated with keratocytes, fibroblasts, and myofibroblasts are differentially expressed between culture conditions. (A) Heatmap of normalized, log2-transformed FPKM values (log2(FPKM+1)) for known keratocyte, fibroblast, and myofibroblast markers. Red color indicates genes with high relative expression levels and blue color indicates genes with lower expression levels. (B, C) Representative western blot and quantification of (B) ALDH1A1 as a marker for keratocytes and (C) Vimentin as a marker for fibroblasts and myofibroblasts. Quantification of target protein expression was normalized to total protein and reported in arbitrary units (a.u.). Error bars represent mean ± s.d. for 3 experimental replicates. A one-way ANOVA with a Tukey post-hoc test was used to compare group means (*, p < 0.05; **, p < 0.01). (D) Confocal overlay images of F-actin, fibronectin, and DAPI staining for corneal keratocytes cultured in SF conditions, or in the presence of FBS or TGF-β1. Scale bar = 20 µm.

Following injury, corneal fibroblasts exhibit an increased proliferative and migratory phenotype, so we also compared levels of expression for genes associated with proliferation and cell-ECM interactions in FBS-treated cells^4, 35^. We observed significant increases in the expression of proliferative markers, such as Ki-67 (MKI67)^36, 37^ and proliferating cell nuclear antigen (PCNA)^38, 39^, as compared to either SF or TGF-β1 (Fig. 2A). In addition, genes encoding vimentin (VIM)^40^, a cytoskeletal intermediate filament, and tenascin C (TNC)^41^, an ECM protein synthesized in repair tissue, also exhibited elevated expression following treatment with FBS. Western blots confirmed increased levels of vimentin in FBS; however, vimentin was also detected at elevated levels in TGF-β1-treated cells (Fig. 2C, S1), a result consistent with previous reports of vimentin expression in myofibroblasts^42^.

We also observed elevated expression of alpha-smooth muscle actin (α-SMA; ACTA2), which incorporates into stress fibers and plays an important role in the increased contractility of myofibroblasts, in both FBS and TGF-β1 as compared to SF (Fig. 2A). In addition, since myofibroblasts have been shown to secrete fibrotic ECM proteins, including biglycan (BGN)^43–46^ and fibronectin (FN1)^47^, we compared the expression levels of BGN and FN1, which were most upregulated upon treatment with TGF-β1. We also observed increased expression of α5 integrin (ITGA5) in both FBS and TGF-β1 (Fig. 2A). Integrin α5β1 is a receptor for fibronectin^48, 49^ and, in previous work, has been shown to be upregulated in both corneal fibroblasts and myofibroblasts^6, 50^. Additionally, consistent with our FN1 expression data, we observed the highest levels of fibronectin immunofluorescence in the presence of TGF-β1 (Fig. 2D), which, as other studies have shown previously^18^, is indicative of a myofibroblast phenotype is this culture condition.

Overall, our RNA-seq data are broadly consistent with known molecular markers for corneal keratocytes, fibroblasts, and myofibroblasts, with SF conditions supporting a keratocyte phenotype, FBS-containing media inducing a fibroblast phenotype, and treatment with TGF-β1 eliciting a myofibroblast phenotype. Even so, genes encoding some fibroblast and myofibroblast markers, such as ITGA5 and ACTA2, were expressed at similar levels in both FBS and TGF-β1 culture conditions, indicating overlap between the gene expression profiles of fibroblasts and myofibroblasts.

### Diverse pathways associated with signal transduction and cellular processes such as cell growth and death, cell-ECM and cell-cell interactions are enriched in the comparisons of fibroblasts and myofibroblasts to keratocytes

KEGG analysis revealed an enrichment of several pathways associated with either signal transduction in TGF-β1-treated myofibroblasts or proliferation in FBS-treated fibroblasts, when compared to keratocytes in SF conditions (Fig. 3). In the presence of FBS (as compared to SF), genes were primarily upregulated in pathways related to translation, replication and repair, and cell growth and death. Among these, we observed significant enrichment of the ribosome and ribosome biogenesis pathways, as well as the DNA replication and cell cycle pathways, suggesting a shift towards increased protein synthesis and cell proliferation upon treatment with FBS (Fig. 3A). In addition, significant changes in expression were observed for genes involved in the PI3K-Akt and Rap1 signaling pathways, as well as those associated with ECM-receptor interactions. These pathways have roles in diverse processes related to proliferation and apoptosis^51–53^, adhesion and migration, and cell-cell and cell-matrix interactions^54–57^.

**Figure 3.**
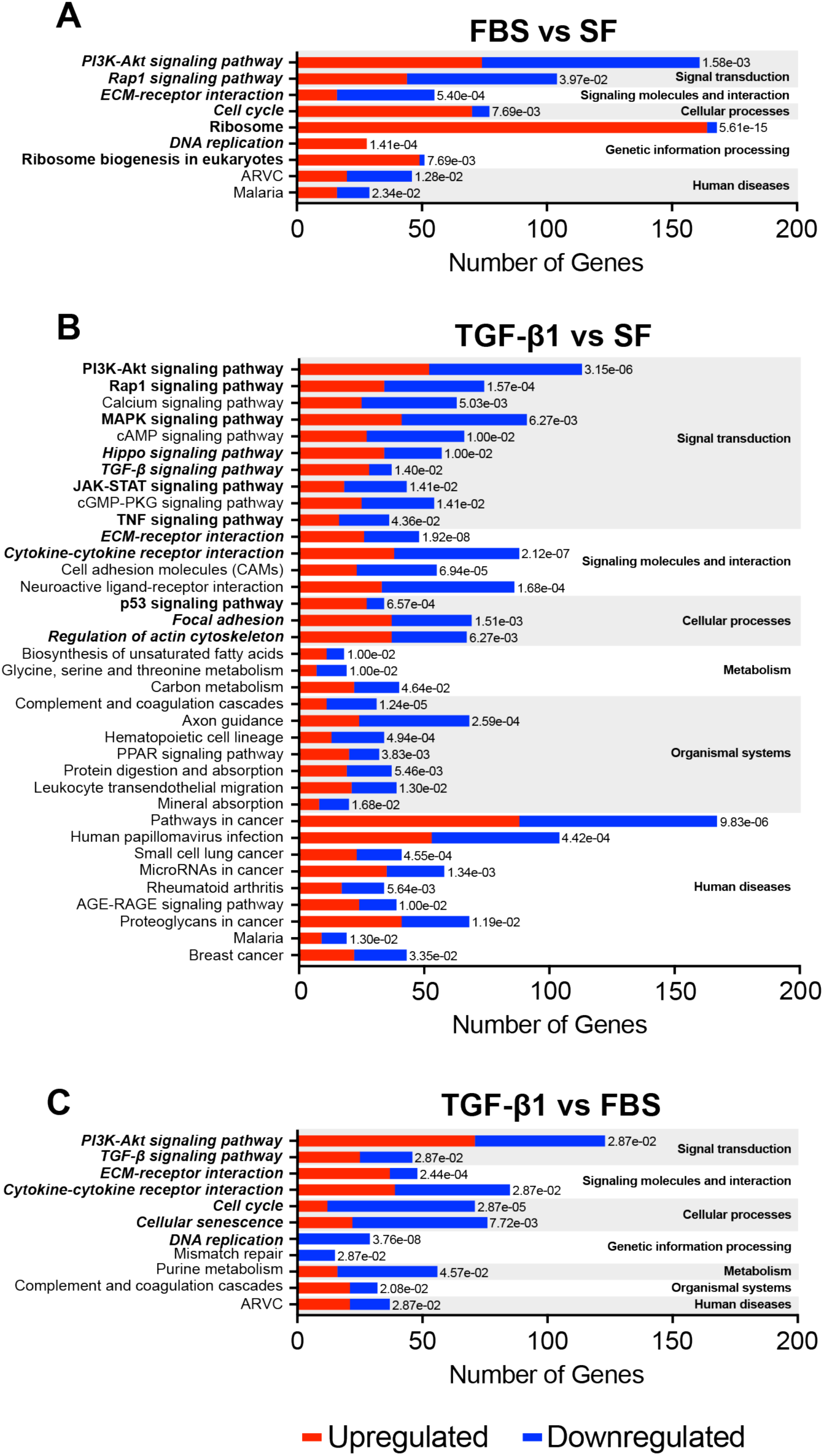
KEGG pathways associated with proliferation are significantly enriched in the comparison of FBS vs. SF, while pathways associated with signal transduction are enriched in the comparison of TGF-β1 vs. SF. Bars represent the number of genes either up- or downregulated in each significantly enriched KEGG pathway from the comparisons of (A) FBS vs. SF, (B) TGF-β1 vs. SF, and (C) TGF-β1 vs. FBS. In each comparison, the alternating grey and white bars differentiate the KEGG pathways grouped together based on their associated function, as indicated on the right of each box. Within each group, pathways have been arranged in order of significance, from most to least significant. KEGG pathways with a padj < 0.05 were considered significant. Pathways of particular interest are highlighted in bold type.

In contrast, the focal adhesion, regulation of actin cytoskeleton, ECM-receptor interaction, and Rap1 signaling KEGG pathways were significantly enriched in the comparison of TGF-β1 to SF, suggesting changes in cytoskeletal organization, cell motility and contractility, and cell-ECM interactions among cultured myofibroblasts (Fig. 3B). Other enriched pathways were broadly related to either cytokine activity (e.g., cytokine-cytokine receptor interaction, as well as TGF-β, JAK-STAT, and TNF signaling pathways) or proliferation and apoptosis (e.g., PI3K-Akt, p53, MAPK, and Hippo signaling pathways).

Strikingly, when TGF-β1-treated myofibroblasts were compared to fibroblasts (as opposed to keratocytes), many of the same KEGG pathways, especially those related to proliferation, cell-ECM interactions, and cytokine activity, were enriched (Fig. 3C). Relative to cells cultured in FBS, TGF-β1 elicited a downregulation of genes associated with the cell cycle and DNA replication pathways, as well as an upregulation of genes in the ECM-receptor interaction pathway. In addition, the cellular senescence pathway, which has been associated with aging, tumor suppression, and wound healing^58, 59^, was also enriched in this comparison. Taken together, these data were suggestive of lower rates of proliferation and a greater emphasis on cell-ECM interactions in TGF-β1-treated myofibroblasts.

### Genes related to proliferation and migration are significantly upregulated in fibroblasts as compared to keratocytes

Numerous genes in the DNA replication, cell cycle, and PI3K-Akt signaling KEGG pathways were differentially expressed in fibroblasts as compared to keratocytes (Fig. 4A-C). Elevated expression was observed for several key genes that encode parts of the DNA replication machinery, including PCNA, MCM2-7, and several DNA polymerase subunits such as POLA2 and POLE2-3^60^. There was also an increase in the expression of several genes associated with the PI3K-Akt signaling pathway, such as the receptor tyrosine kinase MET and the MAP Kinase MAP2K1, which are involved in regulation of proliferation^61, 62^, and several cyclin dependent kinases (CDKs) such as CDK2 and CDK4, which are key regulators of cell cycle progression^63^. The upregulation of these genes was indicative of an increased proliferative phenotype for cultured corneal fibroblasts.

**Figure 4.**
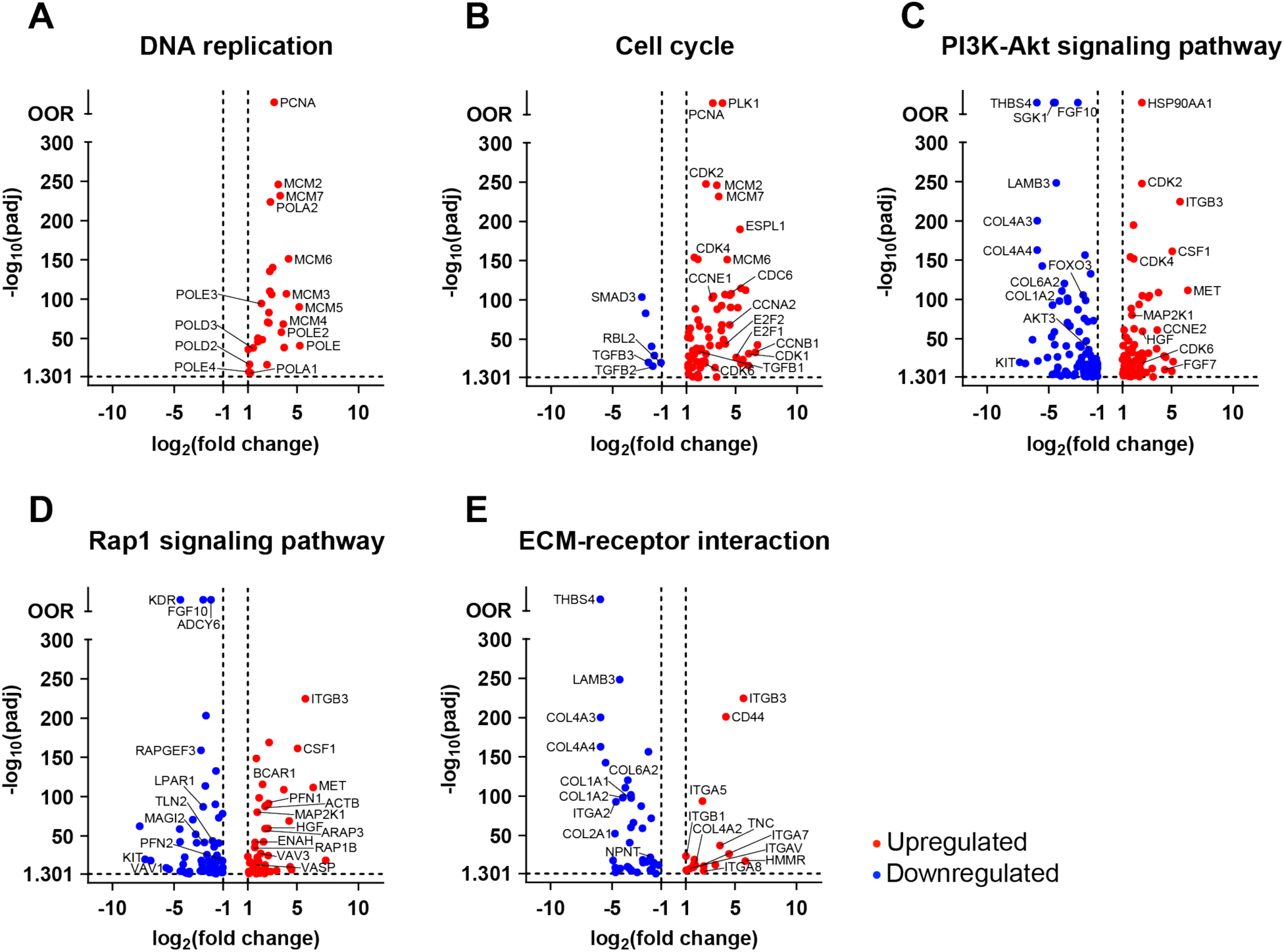
FBS vs. SF: Genes related to a migratory and proliferative phenotype are significantly upregulated in fibroblasts as compared to quiescent keratocytes. (A- E) Volcano plots represent individual genes significantly up- or downregulated within selected KEGG pathways from the comparison of fibroblasts to keratocytes. Genes with padj ≤ 0.05 (horizontal dashed line) and | log2(fold change) | ≥ 1 (vertical dashed lines) were considered significant. Red dots indicate genes that were upregulated in fibroblasts while blue dots indicate downregulated genes. Genes generally associated with FBS induced fibroblast phenotypes and several other significant genes with high padj values or fold changes have been labeled.

Numerous DEGs within the ECM-receptor interaction and Rap1 signaling pathways also indicated a shift toward a migratory phenotype (Fig. 4D-E). Ras-related protein Rap-1b (RAP1B) encodes for a small GTPase that regulates cell adhesion, migration, and polarity^54, 55, 64^. We observed increased expression of RAP1B, along with an upregulation of β-actin (ACTB), which also plays a key role in cell motility^65^. In addition, focal adhesion-related genes, such as talin (TLN2), and genes that regulate actin polymerization, like profilin (PFN2)^66^, were downregulated following treatment with FBS. Changes in cell-ECM interactions were indicated by shifts in the expression of different integrin subunits and many of their corresponding ECM binding partners. This included the downregulation of various collagens (COL1, COL2, COL4, COL6) and the upregulation of tenascin C (TNC), which has been shown to modulate cell adhesion and migration^67, 68^. Consistently, many of the genes encoding integrin subunits that bind to tenascin, such as ITGA8, ITGAV, ITGB1, and ITGB3, were also upregulated. Taken together, these transcriptional changes support an increased proliferative and migratory phenotype in cultured corneal fibroblasts.

### Proliferation related genes are significantly downregulated, while genes involved in cell-ECM interactions and cytokine signaling are upregulated in myofibroblasts compared to fibroblasts

We observed similar differences in proliferation- and motility-related genes when comparing fibroblasts and myofibroblasts. Multiple genes in the DNA replication and cell cycle pathways were differentially expressed between these groups, with lower levels of expression in myofibroblasts indicating a less proliferative phenotype (Fig. 5A-B). Interestingly, genes involved in cell cycle arrest and evasion of apoptosis, such as the cyclin dependent kinase inhibitors 1A and 2B (CDKN1A, CDKN2B), as well as B-cell lymphoma 2 (BCL2), were upregulated in TGF-β1-treated myofibroblasts (Fig. 5C-D)^63, 69–71^.

**Figure 5.**
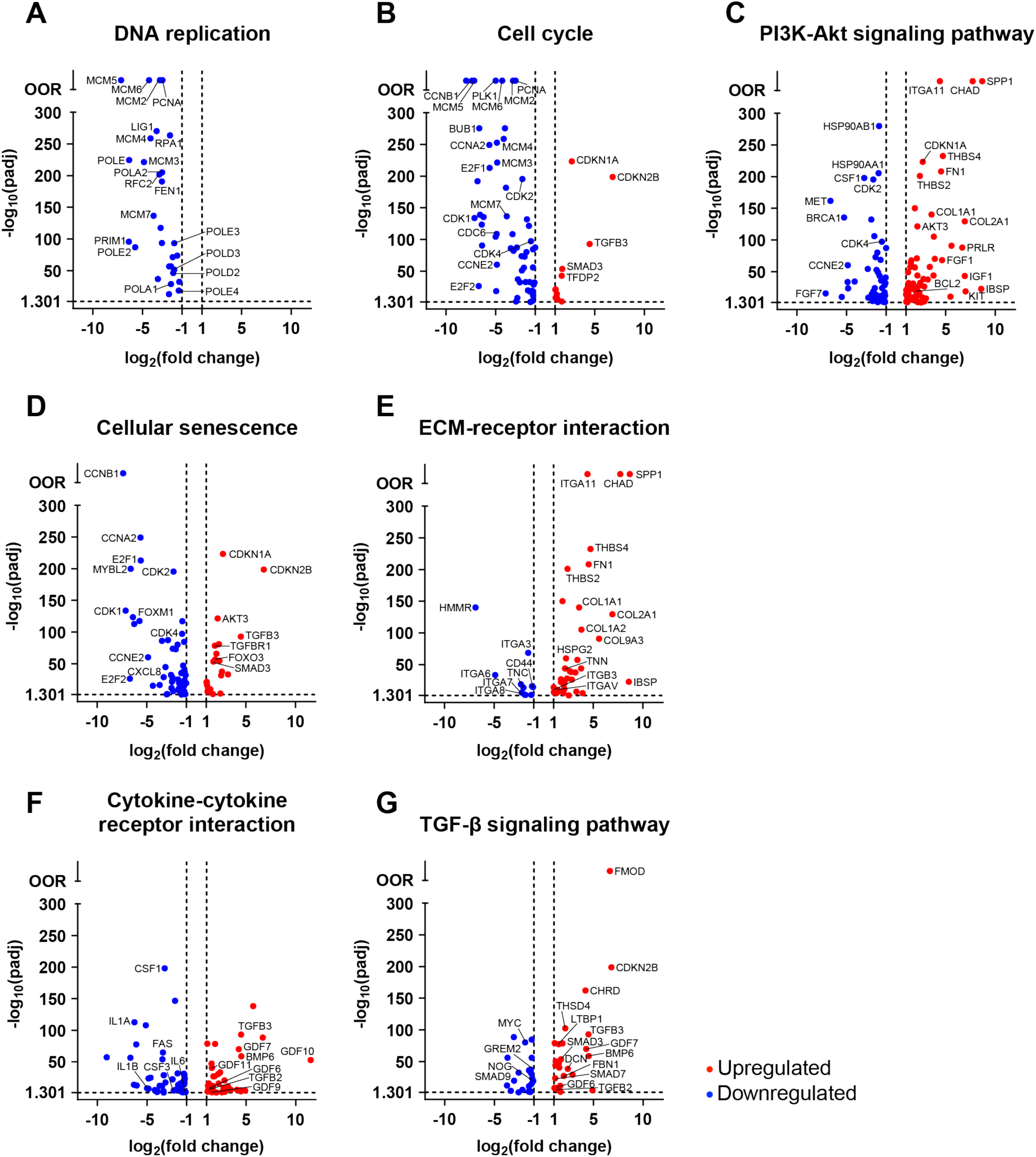
TGF-β1 vs. FBS: Proliferation related genes are significantly downregulated, and genes involved in cell-ECM interactions and cytokine signaling are upregulated in myofibroblasts compared to fibroblasts. (A-G) Volcano plots represent individual genes significantly up- or downregulated within selected KEGG pathways from the comparison of myofibroblasts to fibroblasts. Genes with padj ≤ 0.05 (horizontal dashed line) and | log2(fold change) | ≥ 1 (vertical dashed lines) were considered significant. Red dots indicate genes that were upregulated in myofibroblasts while blue dots indicate downregulated genes. Genes generally associated with either TGF-β1 or FBS induced phenotypes and several other significant genes with high padj values or fold changes have been labeled.

TGF-β1 has been attributed to the induction of ECM synthesis in myofibroblasts^72^. Here, several ECM genes, including fibronectin (FN1), tenascin N (TNN), various collagens (COL1A1, COL1A2, COL2A1, COL9A3), and some of the integrin subunits (e.g., ITGA11, ITGAV, ITGB3) associated with these ECM proteins^57, 73^, exhibited increased expression levels in myofibroblasts compared to fibroblasts (Fig. 5E). Interestingly, the expression of chondroadherin (CHAD), which has been previously shown to promote the adhesion of chondrocytes, fibroblasts, and osteoblasts, was also significantly upregulated in myofibroblasts^74, 75^. In addition, a different gene involved in cell motility – hyaluronan mediated motility receptor (HMMR) – was downregulated in myofibroblasts relative to fibroblasts^76, 77^.

This comparison also revealed significant differences in the expression of genes related to cytokine signaling (Fig. 5F), including (perhaps not surprisingly) several TGF-β family genes (Fig. 5G). Numerous genes encoding TGF-β superfamily ligands, such as growth differentiation factors (GDFs) 6-7 and 9-11, as well as transforming growth factor beta 2 and 3 (TGFB2, TGFB3)^78^, were upregulated in myofibroblasts. We also noted differences in the expression of genes related to cell inflammatory responses, including the interleukins IL1A, IL1B, and IL6, as well as the colony stimulating factors CSF1 and CSF3. These genes were downregulated in myofibroblasts as compared to fibroblasts, which are believed to be the cell type responsible for triggering innate immune responses in the cornea^79^.

In addition, we observed differential expression of several genes associated with TGF-β signaling, some of which encode proteins that are known to modulate TGF-β activity (Fig. 5G). This included the upregulation of two proteoglycans, decorin (DCN) and fibromodulin (FMOD), along with thrombospondin type 1 domain containing 4 (THSD4), fibrillin 1 (FBN1), and latent transforming growth factor beta binding protein 1 (LTBP1)^80^.

### Genes related to cell-ECM interactions and mechanotransduction are differentially expressed in myofibroblasts compared to keratocytes and fibroblasts

Many of the significantly enriched KEGG pathways identified in the comparison of myofibroblasts to keratocytes were associated with cell-ECM interactions and mechanotransduction. These included the ECM-receptor interaction, focal adhesion, regulation of actin cytoskeleton, TGF-β signaling, and Hippo signaling pathways (Fig. 6A-E). Additional KEGG enrichment analysis focused on genes that were significantly upregulated in myofibroblasts revealed significant enrichment of the vascular smooth muscle contraction pathway (Fig. 6F). Consistent with the KEGG analysis, a closer look at the differential gene set for the comparison of myofibroblasts to keratocytes identified many DEGs related to ECM, cell-ECM adhesion, cell-cell interaction, Hippo signaling, actomyosin contractility, and growth factor signaling. Using these categories, genes of interest selected from the highlighted KEGG pathways and the broader differential gene set were classified based on their function and were further evaluated for their relative expression across all three culture conditions (Fig. 7A).

**Figure 6.**
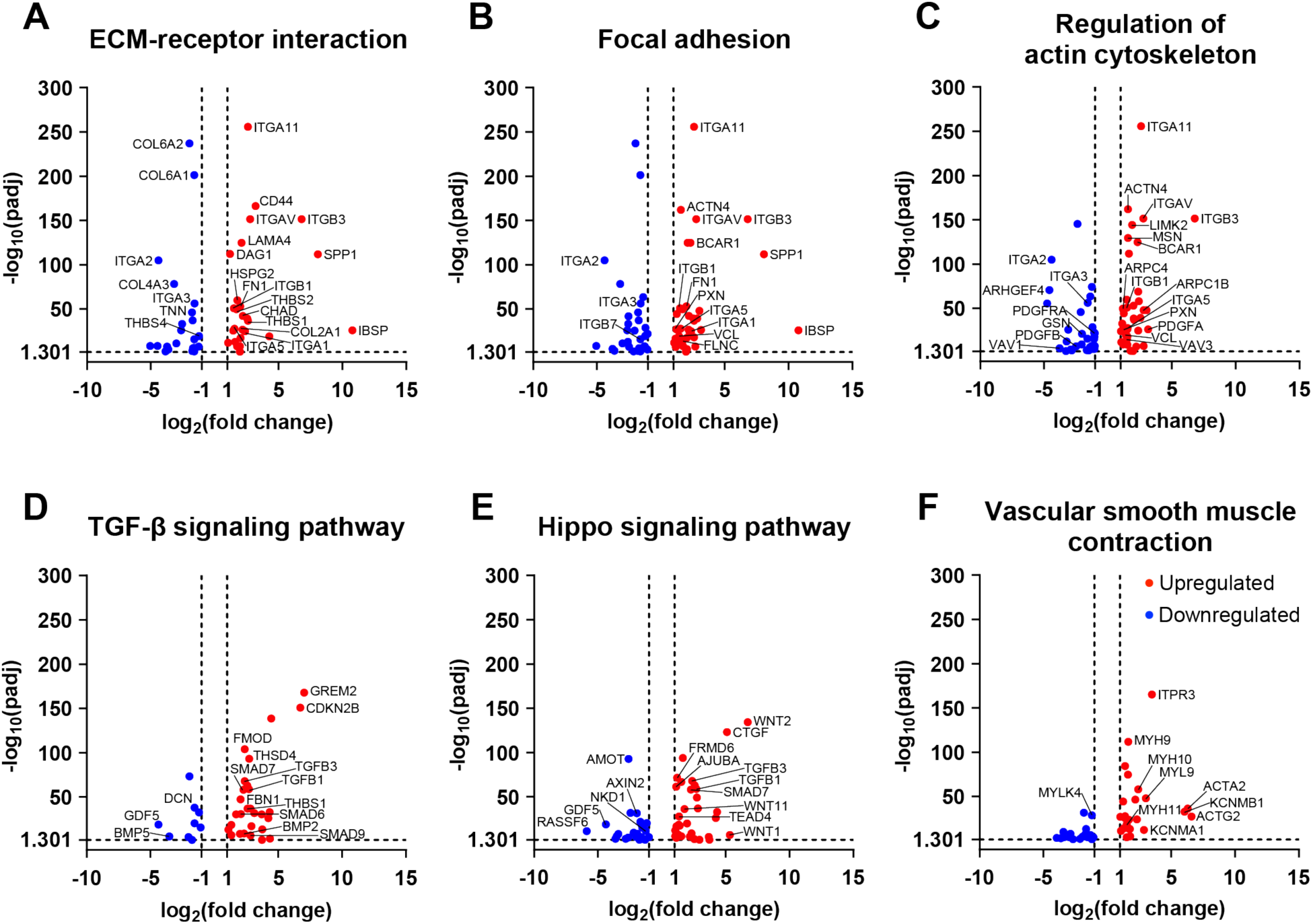
TGF-β1 vs. SF: Genes in KEGG pathways related to cell-ECM interactions and mechanotransduction are differentially expressed in myofibroblasts compared to keratocytes. (A-F) Volcano plots represent individual genes significantly up- or downregulated within selected KEGG pathways from the comparison of myofibroblasts to keratocytes. Genes with padj ≤ 0.05 (horizontal dashed line) and | log2(fold change) | ≥ 1 (vertical dashed lines) were considered significant. Red dots indicate genes that were upregulated in myofibroblasts while blue dots indicate downregulated genes. Genes generally associated with TGF-β1 induced myofibroblast phenotypes and several other significant genes with a high padj values or fold changes have been labeled.

**Figure 7.**
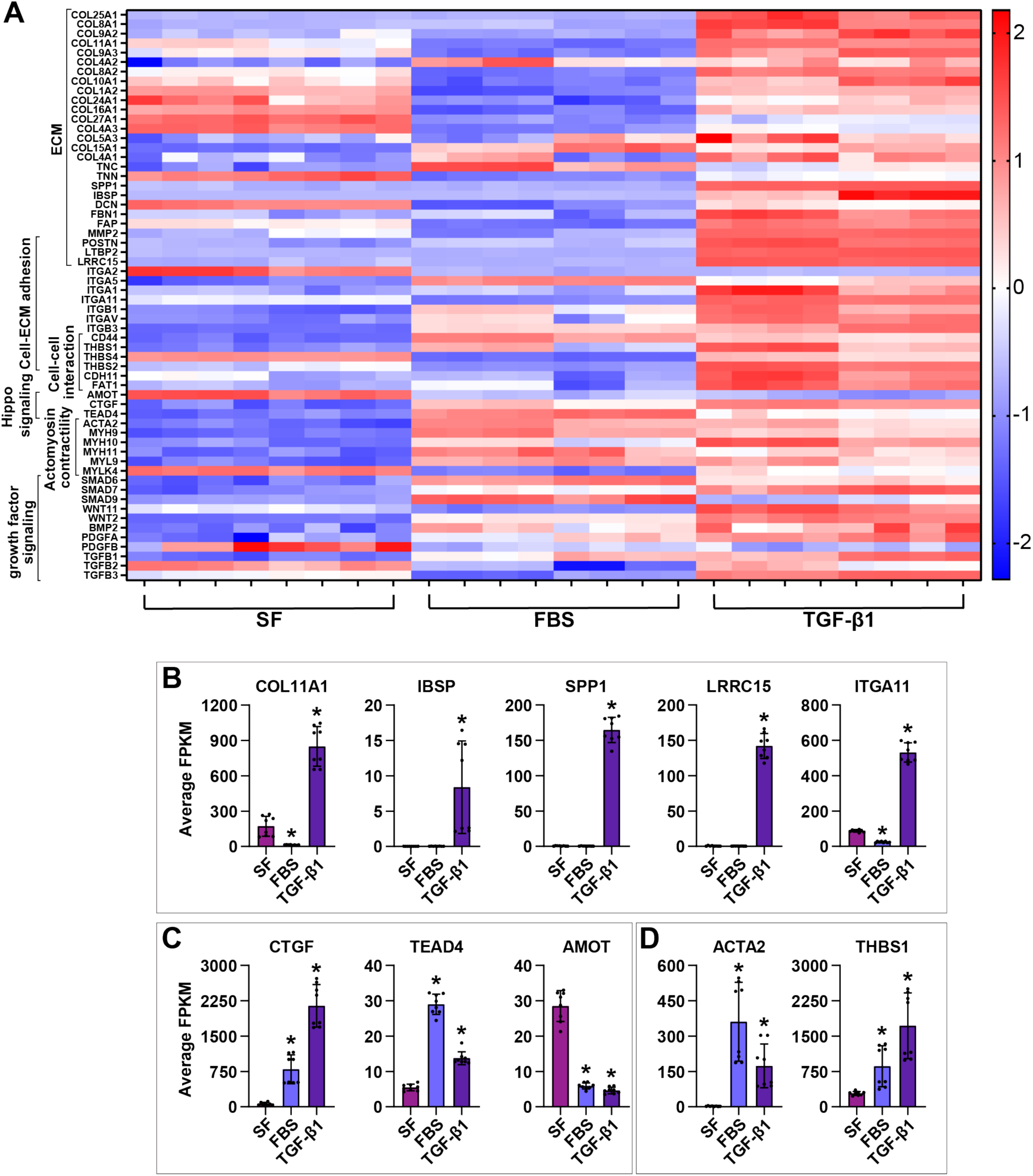
Myofibroblast differentiation is associated with specific ECM and Hippo signaling related genes. (A) Heatmap of normalized, log2-transformed FPKM values (log2(FPKM+1)) for several genes associated with ECM, cell-cell interactions, Hippo signaling, actomyosin contractility, and growth factor signaling that were differentially expressed in myofibroblasts compared to either keratocytes or fibroblasts. Red color indicates genes with high relative expression levels and blue color indicates genes with lower expression levels. Selected genes associated with (B) ECM, (C) Hippo signaling, and (D) myofibroblast differentiation have been highlighted in the bar plots. The average FPKM values have been indicated for comparison among the three culture conditions. Asterisks indicate if expression levels in either FBS or TGF-β1 conditions were found to be significantly different with respect to SF based on the differential expression analysis (padj ≤ 0.05 and | log2(fold change) | ≥ 1).

Multiple collagens were highly expressed in the myofibroblasts compared to keratocytes and fibroblasts. One of the most notable collagens that was significantly upregulated in myofibroblasts was collagen type XI alpha 1 chain (COL11A1) (Fig. 7A, 7B). Interestingly, several other ECM-related genes, such as integrin binding sialoprotein (IBSP), secreted phosphoprotein 1 (SPP1), and leucine rich repeat containing 15 (LRRC15), and integrins such as integrin subunit alpha 11 (ITGA11) were also significantly upregulated in myofibroblasts (Fig. 7A, 7B, Table S1). IBSP is a structural bone matrix protein and SPP1 encodes for osteopontin, known to be important in TGF-β1-induced myofibroblast differentiation^81, 82^. LRRC15 is a membrane protein known to be involved in ECM binding and as a marker for cancer associated myofibroblasts in lung, breast, and other tumors^83–85^. Our data revealed a very significant upregulation of LRRC15 in the presence of TGF-β1, compared to cells cultured in both SF and FBS conditions.

Within the family of genes related to growth factor signaling, a significant upregulation was observed for TGFB1 and TGFB3 in the presence of exogenous TGF-β1 compared to SF (Fig. 7A). Previous work has suggested that TGF-β1 induces the secretion of PDGF as part of an autocrine signaling loop which regulates myofibroblast differentiation^86^. However, it is still unknown which PDGF subunits are involved in this process. Here, our data shows a significant upregulation of PDGFA in myofibroblasts relative to keratocytes and fibroblasts, while PDGFB is downregulated relative to keratocytes (Fig. 7A).

In the comparison of myofibroblasts to keratocytes, DEGs involved in Hippo signaling also stood out due to known crosstalk between this pathway and TGF-β signaling and its association with mechanotransduction^87–89^. Our data revealed differential expression of several Hippo signaling related genes, including connective tissue growth factor (CTGF) and TEA domain transcription factor 4 (TEAD4), which were significantly upregulated in myofibroblasts compared to keratocytes (Fig. 7A, 7C). Interestingly, TEAD4 showed even higher expression in FBS-treated conditions relative to TGF-β1. CTGF, which was most highly expressed in TGF-β1 conditions, is a downstream target of yes-associated protein (YAP) activation, a known mechanosensor^90^. AMOT, encoding for angiomotin, has been known to sequester YAP and TAZ to the cytoplasm, thus reducing the nuclear localization of YAP/TAZ and their downstream effects^91^. Our data indicates that AMOT is significantly downregulated in TGF-β1-treated conditions compared to SF, suggesting nuclear localization of YAP/TAZ.

Several genes involved in actomyosin contractility, such as myosin light/heavy chains and ACTA2, were also upregulated in myofibroblasts as compared to the quiescent keratocytes. ACTA2, a gene encoding for alpha-smooth muscle actin (α-SMA), was significantly upregulated in both fibroblasts and myofibroblasts compared to keratocytes (Fig. 7A, 7D). α-SMA is an important marker for TGF-β1-induced myofibroblast differentiation^25^ and previous studies have shown that the expression of α-SMA in corneal keratocytes is influenced by ECM mechanics^19, 20, 26^.

Another interesting family of genes our data revealed to be differentially expressed in myofibroblasts were the thrombospondins 1,2 and 4 (THBS 1,2,4), known to have the ability to bind to various ECM proteins and play a key role in cell-ECM and cell-cell interactions^92^ (Fig. 7A). THBS1 specifically, was significantly upregulated in myofibroblasts compared to keratocytes (Fig. 7A, 7D) and is known to be important for TGF-β1-induced myofibroblast differentiation^93^.

## Discussion

While many phenotypic differences between corneal keratocytes, fibroblasts, and myofibroblasts have been well documented both *in vivo* and *in vitro*, the transcriptional changes and signaling pathways regulating these differences remain a subject of ongoing research. Improving our understanding of keratocyte differentiation from a genomic perspective could have important implications in corneal wound healing and fibrosis. For this study, we used a 2D *in vitro* model in which corneal keratocytes were cultured in either SF conditions, media containing FBS, or SF media with exogeneous TGF-β1 to produce either corneal keratocytes, fibroblasts, or myofibroblasts, respectively. We then performed bulk RNA sequencing to obtain a global view of the transcriptional differences between each of these cell types. Differential gene expression and functional analyses provided insight into the mechanisms regulating the phenotypic changes associated with each cell type, as well as novel molecular markers and potential targets for modulating cell behavior and differentiation.

Over the years, many previous studies have documented key phenotypic differences between corneal keratocytes, fibroblasts, and myofibroblasts. These include changes in proliferation, ECM synthesis, cell morphology, cytoskeletal organization, and mechanical activity (e.g., adhesion, migration, contractility, etc.)^7, 12, 20, 21, 24–26, 94–96^. Our analyses revealed unique transcriptional profiles for each cell type, which likely underlie these phenotypic differences, since many of the DEGs and enriched signaling pathways were related to proliferation, ECM, and cell mechanics. Among these pathways, we also identified several individual genes that may warrant further investigation to determine their functional significance in the differentiation of keratocytes into fibroblasts and myofibroblasts.

Following injury or surgery, the transformation of quiescent keratocytes into proliferative and migratory fibroblasts is an integral step in the process of corneal wound healing^28, 97^. Our RNA-seq data suggest that the increased proliferation observed among corneal fibroblasts is associated with the upregulation of genes involved in the DNA replication, cell cycle, and PI3K-Akt signaling pathways. This proliferative gene expression signature includes the upregulation of key genes, such as E2F1/2 and CDK2/4/6, in fibroblasts as compared to keratocytes and myofibroblasts. These genes, as well as their associated signaling pathways, may be of further interest as potential targets for accelerating cell repopulation during corneal wound healing or promoting stromal regeneration^98^.

Although the differentiation of myofibroblasts is important for wound closure, a persistent myofibroblast phenotype can lead to protracted corneal fibrosis^97, 99^. Our transcriptional analyses indicated that, although genes associated with proliferation were downregulated, corneal myofibroblasts also appeared to adopt an apoptotic-resistant phenotype associated with pathological wound healing^100, 101^, as indicated by changes in the cell cycle, cellular senescence, PI3K-Akt, and p53 signaling pathways. Genes of interest within these pathways included BCL2, which was upregulated in myofibroblasts compared to both keratocytes and fibroblasts. Due to its role in suppressing apoptosis,

BCL2 has been studied for its therapeutic potential in cancer and during wound healing in other tissues^71, 102, 103^. In general, the evasion of apoptosis by myofibroblasts is considered a hallmark of fibrotic disease, making therapeutic modulation of the cell cycle and apoptosis of interest in a range of pathologies^100, 104, 105^. In the case of corneal wound healing, targeting genes related to apoptosis could be used to support the timely disappearance of myofibroblasts after wound closure to help prevent a fibrotic response.

ECM synthesis and remodeling by corneal keratocytes is important for maintaining the stromal composition and structure that allows for the optical transparency of the tissue. This ECM-maintenance phenotype is disrupted when keratocytes differentiate into fibroblasts and myofibroblasts, a process that changes the types and quantities of ECM proteins produced by the cells^45^. Although previous work has identified several collagens and proteoglycans that are differentially expressed among the different stromal cell types, our data provides a more comprehensive profiling of the many ECM-related genes associated with the matrisome^106, 107^. This includes ECM structural proteins (e.g., collagens, proteoglycans, glycoproteins), cell-ECM receptors (e.g., integrins), ECM-modifying enzymes, and ECM-binding growth factors/cytokines. Altered expression of genes encoding these proteins can produce changes in matrix composition and structure that impact a wide range of cell behaviors, including proliferation, cytoskeletal organization, and mechanical activity^107^.

How the myofibroblast matrisome differs from that of keratocytes and corneal fibroblasts is of particular interest because a key component of the myofibroblast phenotype is the synthesis of an unorganized, fibrotic ECM. Our results are consistent with previous data indicating that the expression of various keratocyte-associated proteoglycans, such as KERA, OGN, and DCN, is reduced in myofibroblasts, while the expression of ECM genes associated with fibrosis (e.g., COL3A1, FN1) is elevated^47^. Our analyses also revealed additional ECM-related genes that were highly upregulated in myofibroblasts and may impact their fibrotic phenotype, including osteopontin (SPP1 or OPN) and integrin binding sialoprotein (IBSP). Each of these genes are primarily expressed within bone matrix, but there is also evidence supporting their role in wound healing and fibrosis in other tissue types^81, 82, 108–112^. OPN expression is important in TGF-β1-induced myofibroblast differentiation within dermal and cardiac fibroblasts and is thought to be a promising therapeutic target for cardiac fibrosis^81, 82^. In addition, OPN knockout mice exhibit delayed corneal wound healing following an incisional injury, which was associated with lower levels of TGF-β1 expression and fewer myofibroblasts^110^.

Changes in ECM composition within the corneal stroma likely also elicit changes in cell behavior via interactions with integrin-containing receptors. Tenascin C (TNC) and fibronectin (FN1), which have been investigated extensively for their involvement in inflammation, wound healing, and tissue remodeling^97, 112, 113^, were found to be highly expressed in corneal fibroblasts and myofibroblasts, respectively. Tenascin C modulates cell adhesion and migration via several integrin receptors, including α8β1 and αvβ1, the subunits of which exhibited elevated gene expression among our cultured fibroblasts. These increases in the expression of TNC and its associated integrin subunits may work together to support the anti-adhesive and pro-migratory phenotypes of corneal fibroblasts, which, along with proliferation, are critical in the early response to corneal injury. In contrast to tenascin C, fibronectin promotes robust cell adhesion to the ECM. Previous work has reported increased fibronectin expression for myofibroblasts, as well as changes in the composition and localization of focal adhesions^20, 21^. Consistent with these data, we observed an upregulation in cultured myofibroblasts of many genes encoding focal adhesion proteins, such as vinculin (VCL), talin (TLN2), and paxillin (PXN). The development of mature focal adhesions is thought to influence the elevated mechanical phenotype associated with myofibroblasts, including increased contractility and traction force generation^21^.

Overall, our data have identified distinct ECM-related gene expression profiles for keratocytes, fibroblasts, and myofibroblasts, which likely contribute to their associated phenotypes. Gene expression by corneal keratocytes promotes a quiescent, ECM-maintenance phenotype, while the altered expression of ECM-related genes promotes a migratory phenotype for fibroblasts and a fibrotic, contractile phenotype for myofibroblasts. How ECM composition and cell-ECM interactions impact corneal wound healing remains a topic of great interest, since ECM structure and composition are important determinants of corneal transparency.

*In vitro* experiments, particularly those employing three-dimensional collagen matrices, have also highlighted the importance of ECM mechanics and stiffness in regulating corneal cell migration, contractility, and ECM reorganization^19, 24, 27, 29, 30^. Furthermore, TGF-β1-induced differentiation of myofibroblasts from quiescent keratocytes is highly sensitive to changes in the mechanical properties of the ECM^20, 24^. In the presence of TGF-β1, keratocytes cultured on softer, uncompressed 3D collagen gels, exhibit fewer stress fibers and reduced expression of α-SMA compared to cells cultured on stiffer, more compressed collagen matrices^14, 19, 94, 95, 114, 115^. Other *in vitro* studies using polyacrylamide gels of varying stiffnesses have shown that corneal keratocytes cultured on stiffer substrata exhibit elevated levels of myofibroblast differentiation in response to TGF-β1, as indicated by an increased number of α-SMA positive cells. This elevated differentiation is also accompanied by changes in both the contractile behavior of the cells and the subcellular localization of focal adhesions^20, 21^.

Given that ECM stiffness is recognized as a significant factor in myofibroblast differentiation, we hypothesized that mechanotransduction-related genes and signaling pathways would be upregulated following culture in the presence of TGF-β1. Indeed, our RNA-seq analyses identified an enrichment of pathways related to cell contractility (e.g., vascular smooth muscle contraction) and mechanotransduction (e.g., Hippo signaling), which include several genes that may contribute to the mechanical differences observed between myofibroblasts and keratocytes. For instance, within the Hippo signaling pathway, we observed changes in the expression of angiomotin (AMOT) and multiple SMADs among myofibroblasts, which are associated with the subcellular localization and activity of YAP/TAZ^91^. Consistent with these data, in previous work, YAP and TAZ have been reported to help regulate myofibroblast differentiation in human corneal fibroblasts^116^. In addition, AMOT has been shown to bind YAP and TAZ, sequester them to the cytoplasm, and thus suppress YAP/TAZ-mediated transcription^91^. In our observation, the downregulation of AMOT among myofibroblasts is consistent with elevated YAP/TAZ activity. This idea is also supported by the increased expression of CTGF, a downstream transcriptional target of YAP activation^90^, in cultured myofibroblasts.

Alpha-smooth muscle actin (α-SMA) expression is a key molecular marker for myofibroblasts. The incorporation of α-SMA into stress fibers is considered a hallmark of myofibroblast differentiation and has been shown to play a significant role in increased contractility and force generation of these cells^13^. Within the vascular smooth muscle contraction pathway, ACTA2, the gene encoding α-SMA, was significantly upregulated in myofibroblasts, as compared to keratocytes. Consistent with previous work, TGF-β1-treated myofibroblasts also contained α-SMA positive stress fibers (Fig. S2A) and western blots showed high α-SMA protein levels relative to keratocytes (Fig. S2B). Interestingly, an even higher level of ACTA2 expression was measured for corneal fibroblasts, although we did not observe the incorporation of α-SMA into stress fibers among cells in this culture condition (Fig. S2A), and western blots showed lower α-SMA protein levels relative to myofibroblasts (Fig. S2B). The RNA-seq data may indicate an important transcriptional difference in ACTA2 expression between corneal fibroblasts and myofibroblasts; however, it is also a possibility that time-varying changes in ACTA2 expression levels contribute to this finding. Furthermore, the western blot data suggest that α-SMA protein levels do not correlate with ACTA2 transcriptional levels when comparing fibroblasts and myofibroblasts, which may indicate potential post-transcriptional or translational control mechanisms.

Taken together, our findings using bulk RNA sequencing provide the first comprehensive differential gene expression profile for cultured corneal keratocytes, fibroblasts, and myofibroblasts. In our initial analysis, we have identified genes and signaling pathways that may play important roles in keratocyte differentiation and the maintenance of these distinct phenotypes. In the future, this sequencing data set can continue to be a resource to identify potential targets for guiding keratocyte differentiation and behavior and serve as a foundation for future studies investigating the influence of substratum stiffness on corneal keratocyte gene expression. Additional experiments targeting specific genes of interest can further reveal the biological and functional significance of these findings as they relate to the broader fields of wound healing, fibrosis, and tissue engineering.

## AUTHOR CONTRIBUTIONS

KP, KSI, VDV, WMP, and DWS conceived the study and designed experiments. KP and KSI conducted all experiments and analyzed all experimental data. All authors discussed and interpreted results. KP, KSI, VDV, and WMP wrote the manuscript with feedback from all authors.

## ACKNOWLEDGMENTS

The authors would also like to thank the members of the Varner, Petroll, and Schmidtke labs for their many helpful discussions and comments.

## Supplemental figures

**Figure S1.**
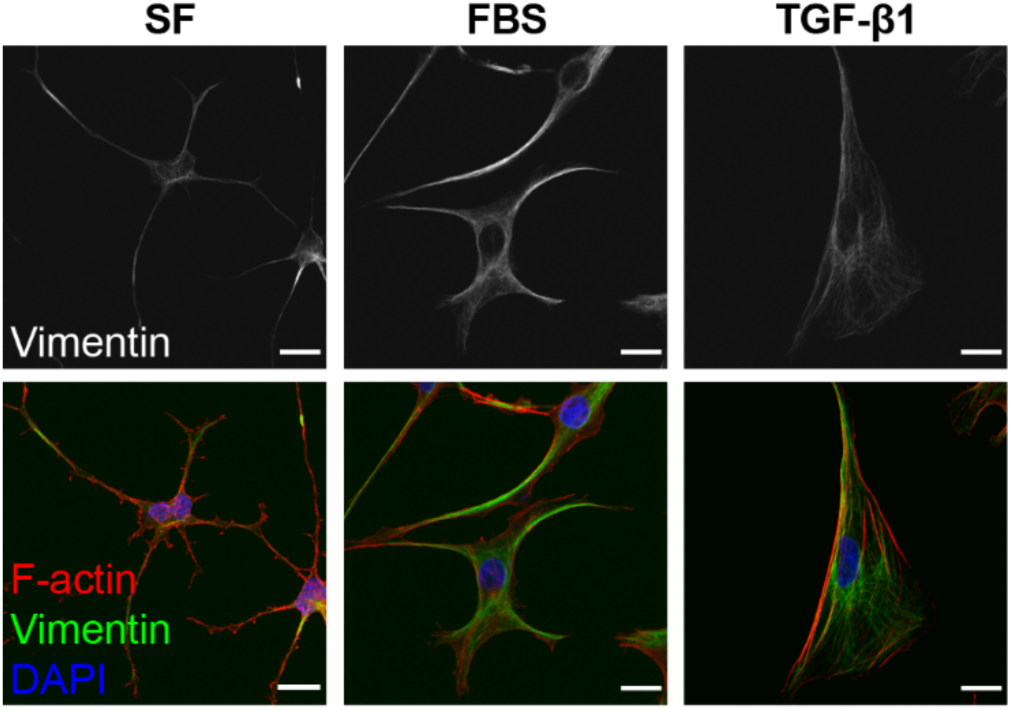
Confocal images of vimentin immunofluorescence and vimentin overlaid with F-actin and DAPI for corneal keratocytes in SF conditions, or in the presence of FBS or TGF-β1. Scale bar = 20 *µ*m.

**Figure S2.**
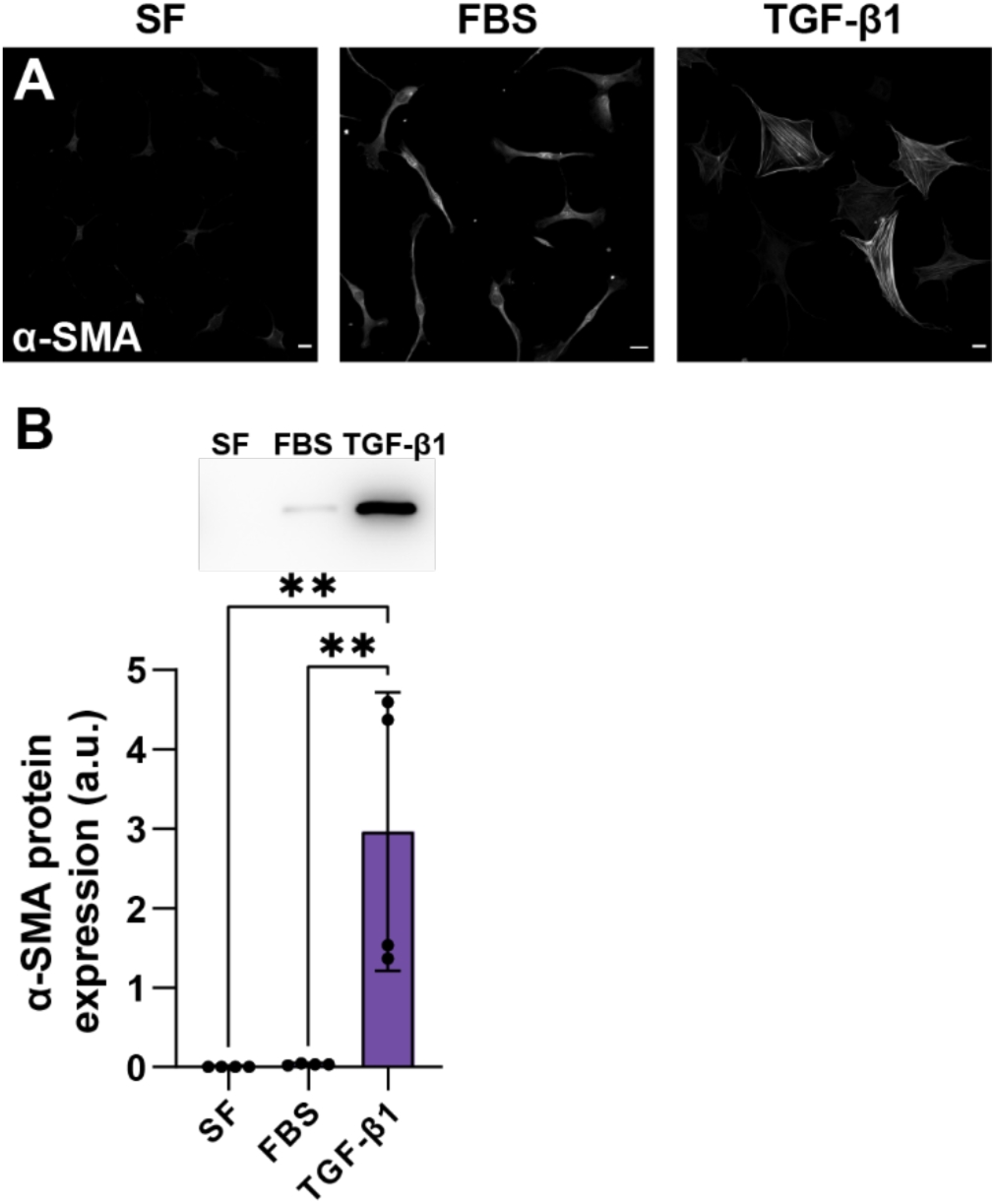
(A) Confocal images of alpha-smooth muscle actin (α-SMA) immunofluorescence for corneal keratocytes in SF, FBS, or TGF-β1 conditions. Scale bar = 20 µm. (B) Representative western blot and quantification of α-SMA expression across culture conditions. Quantification of target protein expression was normalized to total protein and reported in arbitrary units (a.u.). Error bars represent mean ± s.d. for 4 experimental replicates. A one-way ANOVA with a Tukey post-hoc test was used to compare group means (**, p < 0.01).

**Table S1.** Significance values of genes differentially expressed in the presence of FBS or TGF-β1 compared to SF, represented as padj values. List of genes included are those represented as bar plots in Fig. 7B-D.

| | TGF-β1 vs. SF | FBS vs. SF |
| --- | --- | --- |
| GENES | padj | padj |
| COL11A1 | 6.00E-17 | 5.22E-48 |
| IBSP | 9.64E-27 | ns |
| SPP1 | 1.48E-112 | ns |
| LRRC15 | 2.97E-78 | ns |
| ITGA11 | 2.34E-256 | 1.11E-72 |
| CTGF | 9.51E-124 | 3.41E-42 |
| TEAD4 | 1.73E-28 | 3.85E-110 |
| AMOT | 1.45E-93 | 3.34E-79 |
| ACTA2 | 5.52E-37 | 8.79E-53 |
| THBS1 | 1.66E-37 | 6.93E-10 |

## REFERENCES

1. Meek, K.M. and C. Boote, The use of X-ray scattering techniques to quantify the orientation and distribution of collagen in the corneal stroma. Prog Retin Eye Res, 2009. 28(5): p. 369–92.

2. Meek, K.M., Corneal collagen-its role in maintaining corneal shape and transparency. Biophys Rev, 2009. 1(2): p. 83–93.

3. Nishida, T., et al., The network structure of corneal fibroblasts in the rat as revealed by scanning electron microscopy. Invest Ophthalmol Vis Sci, 1988. 29(12): p. 1887–90.

4. Fini, M.E., Keratocyte and fibroblast phenotypes in the repairing cornea. Prog Retin Eye Res, 1999. 18(4): p. 529–51.

5. Worthen, D.M., Histology of the Human Eye. Archives of Ophthalmology, 1972. 88: p. 234–234.

6. Beales, M.P., et al., Proteoglycan Synthesis by Bovine Keratocytes and Corneal Fibroblasts: Maintenance of the Keratocyte Phenotype in Culture. Investigative Ophthalmology & Visual Science, 1999. 40(8): p. 1658–1663.

7. Jester, J.V., et al., Corneal keratocytes: in situ and in vitro organization of cytoskeletal contractile proteins. Invest Ophthalmol Vis Sci, 1994. 35(2): p. 730–43.

8. Zieske, J.D., S.R. Guimarães, and A.E. Hutcheon, Kinetics of keratocyte proliferation in response to epithelial debridement. Exp Eye Res, 2001. 72(1): p. 33–9.

9. Hassell, J.R. and D.E. Birk, The molecular basis of corneal transparency. Exp Eye Res, 2010. 91(3): p. 326–35.

10. Imanishi, J., et al., Growth factors: importance in wound healing and maintenance of transparency of the cornea. Prog Retin Eye Res, 2000. 19(1): p. 113–29.

11. Wilson, S.E., et al., The corneal wound healing response: cytokine-mediated interaction of the epithelium, stroma, and inflammatory cells. Prog Retin Eye Res, 2001. 20(5): p. 625–37.

12. Petroll, W.M., et al., Corneal Fibroblast Migration Patterns During Intrastromal Wound Healing Correlate With ECM Structure and Alignment. Invest Ophthalmol Vis Sci, 2015. 56(12): p. 7352–61.

13. Jester, J.V., et al., Expression of α-smooth muscle (α-SM) actin during corneal stromal wound healing. Investigative Ophthalmology and Visual Science, 1995. 36(5): p. 809–819.

14. Jester, J.V. and J. Ho-Chang, Modulation of cultured corneal keratocyte phenotype by growth factors/cytokines control in vitro contractility and extracellular matrix contraction. Exp Eye Res, 2003. 77(5): p. 581–92.

15. Myrna, K.E., S.A. Pot, and C.J. Murphy, Meet the corneal myofibroblast: the role of myofibroblast transformation in corneal wound healing and pathology. Vet Ophthalmol, 2009. 12 Suppl 1(Suppl 1): p. 25–7.

16. Nakamura, K., Interaction between injured corneal epithelial cells and stromal cells. Cornea, 2003. 22(7 Suppl): p. S35–47.

17. Kurosaka, H., et al., Transforming growth factor-beta 1 promotes contraction of collagen gel by bovine corneal fibroblasts through differentiation of myofibroblasts. Invest Ophthalmol Vis Sci, 1998. 39(5): p. 699–704.

18. Jester, J.V., et al., Transforming Growth Factorβ–Mediated Corneal Myofibroblast Differentiation Requires Actin and Fibronectin Assembly. Investigative Ophthalmology & Visual Science, 1999. 40(9): p. 1959–1967.

19. Petroll, W.M. and M. Miron-Mendoza, Mechanical interactions and crosstalk between corneal keratocytes and the extracellular matrix. Experimental Eye Research, 2015. 133: p. 49–57.

20. Maruri, D.P., et al., ECM Stiffness Controls the Activation and Contractility of Corneal Keratocytes in Response to TGF-β1. Biophys J, 2020. 119(9): p. 1865–1877.

21. Maruri, D.P., et al., Signaling Downstream of Focal Adhesions Regulates Stiffness-Dependent Differences in the TGF-β1-Mediated Myofibroblast Differentiation of Corneal Keratocytes. Frontiers in Cell and Developmental Biology, 2022. 10.

22. Wilson, S.E., Corneal myofibroblast biology and pathobiology: Generation, persistence, and transparency. Experimental Eye Research, 2012. 99: p. 78–88.

23. Jester, J.V., W.M. Petroll, and H.D. Cavanagh, Corneal stromal wound healing in refractive surgery: the role of myofibroblasts. Progress in Retinal and Eye Research, 1999. 18(3): p. 311–356.

24. Kim, A., et al., Growth Factor Regulation of Corneal Keratocyte Differentiation and Migration in Compressed Collagen Matrices. Investigative Ophthalmology & Visual Science, 2010. 51(2): p. 864–875.

25. Jester, J.V., et al., Induction of α-smooth muscle actin expression and myofibroblast transformation in cultured corneal keratocytes. Cornea, 1996. 15(5): p. 505–516.

26. Lakshman, N. and W.M. Petroll, Growth Factor Regulation of Corneal Keratocyte Mechanical Phenotypes in 3-D Collagen Matrices. Investigative Opthalmology & Visual Science, 2012. 53(3): p. 1077.

27. Kim, A., et al., Corneal Stromal Cells use both High- and Low-Contractility Migration Mechanisms in 3-D Collagen Matrices. Experimental cell research, 2012. 318: p. 741–52.

28. Andresen, J.L., T. Ledet, and N. Ehlers, Keratocyte migration and peptide growth factors: the effect of PDGF, bFGF, EGF, IGF-I, aFGF and TGF-beta on human keratocyte migration in a collagen gel. Curr Eye Res, 1997. 16(6): p. 605–13.

29. Kim, A., N. Lakshman, and W.M. Petroll, Quantitative assessment of local collagen matrix remodeling in 3-D Culture: The role of Rho kinase. Experimental Cell Research, 2006. 312(18): p. 3683–3692.

30. Miron-Mendoza, M., et al., The Role of Thrombin and Cell Contractility in Regulating Clustering and Collective Migration of Corneal Fibroblasts in Different ECM Environments. Investigative Ophthalmology & Visual Science, 2015. 56(3): p. 2079–2090.

31. Kivanany, P., et al., An In Vitro Model for Assessing Corneal Keratocyte Spreading and Migration on Aligned Fibrillar Collagen. Journal of Functional Biomaterials, 2018. 9(4): p. 54.

32. Ringnér, M., What is principal component analysis? Nature Biotechnology, 2008. 26(3): p. 303–304.

33. Hassell, J.R., et al., Proteoglycan changes during restoration of transparency in corneal scars. Archives of Biochemistry and Biophysics, 1983. 222(2): p. 362–369.

34. Jester, J.V., Corneal crystallins and the development of cellular transparency. Semin Cell Dev Biol, 2008. 19(2): p. 82–93.

35. Mohan, R.R., et al., Apoptosis, necrosis, proliferation, and myofibroblast generation in the stroma following LASIK and PRK. Exp Eye Res, 2003. 76(1): p. 71–87.

36. McKay, T.B., et al., Extracellular Vesicles Secreted by Corneal Epithelial Cells Promote Myofibroblast Differentiation. Cells, 2020. 9(5): p. 1080.

37. Zixian, D., et al., Small incision lenticule extraction (SMILE) and femtosecond laser LASIK: comparison of corneal wound healing and inflammation. British Journal of Ophthalmology, 2014. 98(2): p. 263.

38. binte M. Yusoff, N.Z., et al., Isolation and Propagation of Human Corneal Stromal Keratocytes for Tissue Engineering and Cell Therapy. Cells, 2022. 11(1): p. 178.

39. Gan, L., P. Fagerholm, and S. Ekenbark, Expression of proliferating cell nuclear antigen in corneas kept in long term culture. Acta Ophthalmol Scand, 1998. 76(3): p. 308–13.

40. Wilson, S.E., The Yin and Yang of Mesenchymal Cells in the Corneal Stromal Fibrosis Response to Injury: The Cornea as a Model of Fibrosis in Other Organs. Biomolecules, 2022. 13(1).

41. Latvala, T., et al., Expression of cellular fibronectin and tenascin in the rabbit cornea after excimer laser photorefractive keratectomy: a 12 month study. British Journal of Ophthalmology, 1995. 79(1): p. 65.

42. Chaurasia, S.S., et al., Dynamics of the expression of intermediate filaments vimentin and desmin during myofibroblast differentiation after corneal injury. Experimental Eye Research, 2009. 89(2): p. 133–139.

43. Funderburgh, J.L., et al., Proteoglycan expression during transforming growth factor beta -induced keratocyte-myofibroblast transdifferentiation. J Biol Chem, 2001. 276(47): p. 44173–8.

44. Funderburgh, J.L., et al., Decorin and biglycan of normal and pathologic human corneas. Investigative Ophthalmology & Visual Science, 1998. 39(10): p. 1957–1964.

45. Funderburgh, J.L., M.M. Mann, and M.L. Funderburgh, Keratocyte phenotype mediates proteoglycan structure: a role for fibroblasts in corneal fibrosis. J Biol Chem, 2003. 278(46): p. 45629–37.

46. Massoudi, D., F. Malecaze, and S.D. Galiacy, Collagens and proteoglycans of the cornea: importance in transparency and visual disorders. Cell and Tissue Research, 2016. 363(2): p. 337–349.

47. Garana, R.M.R., et al., Radial keratotomy: II. Role of the myofibroblast in corneal wound contraction. Investigative Ophthalmology and Visual Science, 1992. 33(12): p. 3271–3282.

48. Welch, M.P., G.F. Odland, and R.A. Clark, Temporal relationships of F-actin bundle formation, collagen and fibronectin matrix assembly, and fibronectin receptor expression to wound contraction. J Cell Biol, 1990. 110(1): p. 133–45.

49. Schumacher, S., et al., Structural insights into integrin α(5)β(1) opening by fibronectin ligand. Sci Adv, 2021. 7(19).

50. Masur, S.K., J.K. Cheung, and S. Antohi, Identification of integrins in cultured corneal fibroblasts and in isolated keratocytes. Investigative Ophthalmology & Visual Science, 1993. 34(9): p. 2690–2698.

51. Engelman, J.A., J. Luo, and L.C. Cantley, The evolution of phosphatidylinositol 3-kinases as regulators of growth and metabolism. Nat Rev Genet, 2006. 7(8): p. 606–19.

52. Kim, D. and J. Chung, Akt: versatile mediator of cell survival and beyond. J Biochem Mol Biol, 2002. 35(1): p. 106–15.

53. Chen, K., et al., The role of the PI3K/AKT signalling pathway in the corneal epithelium: recent updates. Cell Death Dis, 2022. 13(5): p. 513.

54. Bos, J.L., Linking Rap to cell adhesion. Curr Opin Cell Biol, 2005. 17(2): p. 123–8.

55. Boettner, B. and L. Van Aelst, Control of cell adhesion dynamics by Rap1 signaling. Curr Opin Cell Biol, 2009. 21(5): p. 684–93.

56. Bosman, F.T. and I. Stamenkovic, Functional structure and composition of the extracellular matrix. J Pathol, 2003. 200(4): p. 423–8.

57. van der Flier, A. and A. Sonnenberg, Function and interactions of integrins. Cell Tissue Res, 2001. 305(3): p. 285–98.

58. Childs, B.G., et al., Cellular senescence in aging and age-related disease: from mechanisms to therapy. Nat Med, 2015. 21(12): p. 1424–35.

59. Muñoz-Espín, D. and M. Serrano, Cellular senescence: from physiology to pathology. Nat Rev Mol Cell Biol, 2014. 15(7): p. 482–96.

60. Waga, S. and B. Stillman, The DNA replication fork in eukaryotic cells. Annu Rev Biochem, 1998. 67: p. 721–51.

61. Trusolino, L., A. Bertotti, and P.M. Comoglio, MET signalling: principles and functions in development, organ regeneration and cancer. Nat Rev Mol Cell Biol, 2010. 11(12): p. 834–48.

62. Organ, S.L. and M.S. Tsao, An overview of the c-MET signaling pathway. Ther Adv Med Oncol, 2011. 3(1 Suppl): p. S7–s19.

63. Łukasik, P., M. Załuski, and I. Gutowska, Cyclin-Dependent Kinases (CDK) and Their Role in Diseases Development-Review. Int J Mol Sci, 2021. 22(6).

64. Wilson, J.M., et al., Differences in the Phosphorylation-Dependent Regulation of Prenylation of Rap1A and Rap1B. J Mol Biol, 2016. 428(24 Pt B): p. 4929–4945.

65. Bunnell, T.M., et al., β-Actin specifically controls cell growth, migration, and the G-actin pool. Mol Biol Cell, 2011. 22(21): p. 4047–58.

66. Ding, Z. and P. Roy, Profilin-1 versus profilin-2: two faces of the same coin? Breast Cancer Res, 2013. 15(3): p. 311.

67. Jones, F.S. and P.L. Jones, The tenascin family of ECM glycoproteins: structure, function, and regulation during embryonic development and tissue remodeling. Dev Dyn, 2000. 218(2): p. 235–59.

68. Midwood, K.S. and G. Orend, The role of tenascin-C in tissue injury and tumorigenesis. J Cell Commun Signal, 2009. 3(3-4): p. 287–310.

69. Bartkova, J., et al., Cell-cycle regulatory proteins in human wound healing. Archives of Oral Biology, 2003. 48(2): p. 125–132.

70. Reed, J.C., Bcl-2 and the regulation of programmed cell death. J Cell Biol, 1994. 124(1-2): p. 1–6.

71. Ruvolo, P.P., X. Deng, and W.S. May, Phosphorylation of Bcl2 and regulation of apoptosis. Leukemia, 2001. 15(4): p. 515–22.

72. Wilson, S.E., Corneal myofibroblasts and fibrosis. Exp Eye Res, 2020. 201: p. 108272.

73. Degen, M., et al., Tenascin-W is a novel marker for activated tumor stroma in low-grade human breast cancer and influences cell behavior. Cancer Res, 2007. 67(19): p. 9169–79.

74. Mizuno, M., R. Fujisawa, and Y. Kuboki, Bone chondroadherin promotes attachment of osteoblastic cells to solid-state substrates and shows affinity to collagen. Calcif Tissue Int, 1996. 59(3): p. 163–7.

75. Camper, L., D. Heinegârd, and E. Lundgren-Akerlund, Integrin alpha2beta1 is a receptor for the cartilage matrix protein chondroadherin. J Cell Biol, 1997. 138(5): p. 1159–67.

76. Tolg, C., et al., Hyaluronan and RHAMM in wound repair and the “cancerization” of stromal tissues. Biomed Res Int, 2014. 2014: p. 103923.

77. Hinneh, J.A., et al., The role of RHAMM in cancer: Exposing novel therapeutic vulnerabilities. Front Oncol, 2022. 12: p. 982231.

78. Wrana, J.L., Signaling by the TGFβ superfamily. Cold Spring Harb Perspect Biol, 2013. 5(10): p. a011197.

79. Fukuda, K., Corneal fibroblasts: Function and markers. Experimental Eye Research, 2020. 200: p. 108229.

80. Soo, C., et al., Differential expression of fibromodulin, a transforming growth factor-beta modulator, in fetal skin development and scarless repair. Am J Pathol, 2000. 157(2): p. 423–33.

81. Lenga, Y., et al., Osteopontin expression is required for myofibroblast differentiation. Circ Res, 2008. 102(3): p. 319–27.

82. Abdelaziz Mohamed, I., et al., Osteopontin: A Promising Therapeutic Target in Cardiac Fibrosis. Cells, 2019. 8(12).

83. Purcell, J.W., et al., LRRC15 Is a Novel Mesenchymal Protein and Stromal Target for Antibody-Drug Conjugates. Cancer Res, 2018. 78(14): p. 4059–4072.

84. Krishnamurty, A.T., et al., LRRC15(+) myofibroblasts dictate the stromal setpoint to suppress tumour immunity. Nature, 2022. 611(7934): p. 148–154.

85. Krishnamurty, A., et al., 1434 TGFβ-dependent LRRC15^+^ myofibroblasts dictate the tumor fibroblast setpoint to promote cancer immunotherapy resistance. Journal for ImmunoTherapy of Cancer, 2023. 11(Suppl 1): p. A1597–A1597.

86. Jester, J.V., et al., TGFβ Induced Myofibroblast Differentiation of Rabbit Keratocytes Requires Synergistic TGFβ, PDGF and Integrin Signaling. Experimental Eye Research, 2002. 75(6): p. 645–657.

87. Attisano, L. and J.L. Wrana, Signal integration in TGF-β, WNT, and Hippo pathways. F1000Prime Rep, 2013. 5: p. 17.

88. Chang, Y.C., et al., Hippo Signaling-Mediated Mechanotransduction in Cell Movement and Cancer Metastasis. Front Mol Biosci, 2019. 6: p. 157.

89. Cobbaut, M., et al., Dysfunctional Mechanotransduction through the YAP/TAZ/Hippo Pathway as a Feature of Chronic Disease. Cells, 2020. 9(1): p. 151.

90. Raghunathan, V.K., et al., Involvement of YAP, TAZ and HSP90 in contact guidance and intercellular junction formation in corneal epithelial cells. PLoS One, 2014. 9(10): p. e109811.

91. Chan, S.W., et al., Hippo pathway-independent restriction of TAZ and YAP by angiomotin. J Biol Chem, 2011. 286(9): p. 7018–26.

92. Lahav, J., The functions of thrombospondin and its involvement in physiology and pathophysiology. Biochim Biophys Acta, 1993. 1182(1): p. 1–14.

93. Bradshaw, A.D., *Chapter 15 - Regulation of Cell Behavior by Extracellular Proteins*, in Principles of Tissue Engineering (Fourth Edition), R. Lanza, R. Langer, and J. Vacanti, Editors. 2014, Academic Press: Boston. p. 279–290.

94. Lakshman, N., A. Kim, and W.M. Petroll, Characterization of corneal keratocyte morphology and mechanical activity within 3-D collagen matrices. Experimental Eye Research, 2010. 90(2): p. 350–359.

95. Karamichos, D., N. Lakshman, and W.M. Petroll, Regulation of Corneal Fibroblast Morphology and Collagen Reorganization by Extracellular Matrix Mechanical Properties. Investigative Ophthalmology & Visual Science, 2007. 48(11): p. 5030–5037.

96. Iyer, K.S., et al., ECM stiffness modulates the proliferation but not the motility of primary corneal keratocytes in response to PDGF-BB. Experimental Eye Research, 2022. 220: p. 109112.

97. Kamil, S. and R.R. Mohan, Corneal stromal wound healing: Major regulators and therapeutic targets. Ocul Surf, 2021. 19: p. 290–306.

98. Lagali, N., Corneal Stromal Regeneration: Current Status and Future Therapeutic Potential. Curr Eye Res, 2020. 45(3): p. 278–290.

99. Desai, V.D., H.C. Hsia, and J.E. Schwarzbauer, Reversible Modulation of Myofibroblast Differentiation in Adipose-Derived Mesenchymal Stem Cells. PLOS ONE, 2014. 9(1): p. e86865.

100. Hinz, B. and D. Lagares, Evasion of apoptosis by myofibroblasts: a hallmark of fibrotic diseases. Nat Rev Rheumatol, 2020. 16(1): p. 11–31.

101. McElhinney, K., M. Irnaten, and C. O’Brien, p53 and Myofibroblast Apoptosis in Organ Fibrosis. International Journal of Molecular Sciences, 2023. 24(7): p. 6737.

102. Qian, S., et al., The role of BCL-2 family proteins in regulating apoptosis and cancer therapy. Front Oncol, 2022. 12: p. 985363.

103. Wilkinson, H.N. and M.J. Hardman, Senescence in Wound Repair: Emerging Strategies to Target Chronic Healing Wounds. Front Cell Dev Biol, 2020. 8: p. 773.

104. Wu, Y.S., et al., Cell Cycle Dysregulation and Renal Fibrosis. Front Cell Dev Biol, 2021. 9: p. 714320.

105. Rosenwald, A., et al., The proliferation gene expression signature is a quantitative integrator of oncogenic events that predicts survival in mantle cell lymphoma. Cancer Cell, 2003. 3(2): p. 185–97.

106. Naba, A., et al., The extracellular matrix: Tools and insights for the “omics” era. Matrix Biology, 2016. 49: p. 10–24.

107. Hynes, R.O. and A. Naba, Overview of the matrisome--an inventory of extracellular matrix constituents and functions. Cold Spring Harb Perspect Biol, 2012. 4(1): p. a004903.

108. Icer, M.A. and M. Gezmen-Karadag, The multiple functions and mechanisms of osteopontin. Clin Biochem, 2018. 59: p. 17–24.

109. Sumioka, T., et al., Tenascins and osteopontin in biological response in cornea. Ocul Surf, 2023. 29: p. 131–149.

110. Miyazaki, K.-i., et al., Corneal Wound Healing in an Osteopontin-Deficient Mouse. Investigative Ophthalmology & Visual Science, 2008. 49(4): p. 1367–1375.

111. Moorman, H.R., et al., Osteopontin: A Key Regulator of Tumor Progression and Immunomodulation. Cancers, 2020. 12(11): p. 3379.

112. Saika, S., et al., Modulation of Smad signaling by non-TGFβ components in myofibroblast generation during wound healing in corneal stroma. Exp Eye Res, 2016. 142: p. 40–8.

113. Varkoly, G., et al., Expression Pattern of Tenascin-C, Matrilin-2, and Aggrecan in Diseases Affecting the Corneal Endothelium. Journal of Clinical Medicine, 2022. 11(20): p. 5991.

114. Raghunathan, V.K., et al., Tissue and cellular biomechanics during corneal wound injury and repair. Acta Biomaterialia, 2017. 58: p. 291–301.

115. Dreier, B., et al., Substratum Compliance Modulates Corneal Fibroblast to Myofibroblast Transformation. Investigative Ophthalmology & Visual Science, 2013. 54(8): p. 5901–5907.

116. Muppala, S., et al., YAP and TAZ are distinct effectors of corneal myofibroblast transformation. Exp Eye Res, 2019. 180: p. 102–109.

